# Large scale capsid-mediated mobilisation of bacterial genomic DNA in the gut microbiome

**DOI:** 10.1101/2024.11.15.623857

**Authors:** Tatiana Borodovich, Jason S. Wilson, Pavol Bardy, Muireann Smith, Conor Hill, Ekaterina V. Khokhlova, Bianca Govi, Paul C.M. Fogg, Colin Hill, Andrey N. Shkoporov

## Abstract

Transducing bacteriophages and gene transfer agents (GTA) are constrained by their capsids’ structural properties in the length of host DNA they can package. Nanopore sequencing of intact capsid-packaged DNA molecules with full-sized reads can be used to establish the precise lengths and identity of individual packaged DNA molecules and their association with specific bacterial hosts. This approach was validated using a few well-characterised transducing systems, and then applied to study bacterial DNA encapsidation in the faecal microbiomes from three healthy human donors. Bacterial DNA encapsidation appears to be widespread in the microbiome with up to 5.4% of capsid-packaged DNA in the gut virome being of bacterial (non-prophage) origin. Generalised transduction and GTA activity are especially prevalent in the families Oscillospiraceae and Ruminococcaceae, whereas an example of lateral transduction was observed in genus Bacteroides. In addition to that, induction of prophages in a variety of highly prevalent gut bacteria was observed.

## Introduction

High taxonomic diversity^1^ and functional redundancy^2^ are essential properties of the human gut microbiome and other complex microbial communities. These properties are believed to underpin the resilience of the microbiome – its ability to rebound to a functionally similar (if not the same) state after removal of a stressing factor causing its major perturbation, such as a sudden change in diet, a disease state affecting the gut environment, or antibiotic therapy^3,4^. Recent evolutionary modelling proposed that horizontal gene transfer (HGT; potentially including phage-mediated HGT) can help to overcome diversity limits for competing microbial species and strains through sharing of fitness determinants and establishment of “dynamic neutrality”^5^. In line with this, empirical data from large-scale gut microbial genomics and metagenomics studies highlight extensive recent horizontal acquisitions in bacterial genomes^6^. HGT is especially common in the industrialised population microbiomes^7^, with at least some of these events linked with phage transduction.

Transduction, in general, can refer to any HGT mechanism in which DNA is encapsulated inside a nanoparticle, such as a proteinaceous capsid or lipid vesicle etc, and transferred from a donor to a recipient cell. Herein, we will refer to transduction specifically as a transfer of DNA within a virus or a virus-like capsid^8^. Specialized phage transduction (ST) occurs when a fragment of the bacterial genome immediately adjacent to the prophage is occasionally packaged together with the phage DNA. Generalized transduction (GT) can package consecutive DNA fragments covering extensive regions of bacterial genomes, if not entire genomes, and is typically initiated by recognition of pseudo-*pac* sites within the bacterial chromosome by a sequence-specific phage terminase. The more recently discovered lateral transduction (LT) is driven by prophage replication inside the host chromosome prior to excision, and results in packaging of large spans of host DNA, in consecutive fragments dictated by the phage “headful” DNA capacity (capsid capacity), extending from one end of the integrated prophage genome^9^. Temperate phages and phage-inducible chromosomal islands (PICI) capable of LT have been described in *Staphylococcus aureus*^9–11^, *Enterococcus faecalis*^12^ and *Salmonella enterica*^13,14^. Lateral transduction in *S. aureus* is characterised by a very high efficiency^9,10^.

Gene transfer agents (GTAs) are small phage-like particles that have independently arisen in multiple bacterial lineages through “domestication” of ancestral phages^15,16^. All characterized GTAs are encoded by several operons spread across the host bacterial genome and are incapable of autonomous replication^17^. The DNA packaging motor is comparable to phage terminase complexes, however the terminase lacks DNA sequence specificity and can potentially process any DNA within the cell^18^. Only a handful of GTAs have been characterized in-depth to date, mainly in Alphaproteobacteria such as *Rhodobacter*, *Dinoroseobacter, Bartonella* and *Caulobacter*^19^.

Bioinformatic discovery of new GTA-like elements is a non-trivial task, due to their similarity to defective prophages and degraded prophage remnants. Furthermore, unlike transducing temperate phages, known GTA elements described in *Rhodobacterales*^20–22^, *Desulfovibrio*^23^, *Brachyspira*^24^, *Bartonella spp.*^25^, *Caulobacter*^26^, and *Methanococcus voltae*^27^, do not engage in preferential packaging of the elements themselves, instead processing and packaging the entire chromosome in a (quasi-)non-specific manner. The lack of selection for self-DNA inside GTA particles is another obstacle to identification of the GTA encoding genes.

Capturing ongoing transduction events in the gut microbiome and other similarly complex microbial environments remains a challenge, especially between members of a single species. Observations of significant quantities of bacterial DNA associated with virus-like particles (VLP) in gut virome studies^28^ have often been ascribed to contamination^29^. When short-read shotgun sequencing is used and no information on the size or topology of VLP-associated DNA molecules is available, contaminating DNA released from bacterial cells may be indistinguishable from the contents of true transducing particles. This is further compounded by possible co-purification of membrane vesicles along with VLPs, with some of them capable of carrying DNA (vesicle-mediated transduction)^30–32^. A pioneering approach to address this problem (termed “transductomics”), consists in mapping of the VLP DNA reads to metagenomically assembled bacterial contigs and genomes from the same microbiome, to reveal coverage patterns consistent with known phage transduction mechanisms^12^. This approach, however, suffers from low resolution, as short DNA reads are treated as a collective, rather than on individual basis, and some of the patterns produced can be explained by more than one, or none of the known transduction mechanisms.

The size of phage-encapsidated DNA is determined by the packaging mechanism and capsid capacity^33,34^. Tailed phages containing cohesive DNA ends (*cos* sites in *E. coli* phage phage λ), or double stranded terminal direct repeats (short in *E. coli* phage T7, long in T5) package identical copies of DNA of precise length. “Headful” packaging of a concatemeric DNA precursor (initiated from a *pac* site in *Salmonella* phage P22 and *E. coli* phage P1) results in terminally redundant, circularly permuted fragments, each being 2%-10% longer than the length of the unique genome sequence. Headful packaging machinery typically allows for ±2% imprecision of the packaged DNA size. Phage Mu (and related phages) combines replicative transposition with a somewhat unique headful packaging mechanism, resulting in packaging of 500-3,000 bp of variable host DNA in addition to its own 37 kbp genome. Therefore, regardless of the type of the specific phage transduction mechanism, the length of packaged DNA is determined by the capsid size and specific packaging mechanism and remains relatively constant for each capsid. Similarly, transducing particles (defective virions) formed by tailed phages capable of GT and LT are expected to contain bacterial DNA fragments of size similar to the size of packaged phage DNA in a normal virion. In contrast, GTAs from *Alphaproteobacteria* have been shown to package DNA with ∼20% reduced density into an oblate T=3 capsid, explaining the ∼4kb capacity of the capsid^35^.

Here we propose, that extracting minimally damaged DNA from faecal VLP fractions and subjecting it to direct Nanopore sequencing with terminally ligated sequencing adaptors makes it possible to capture the intact DNA content of individual virions (both normal virions and transducing particles) and build a holistic view of ongoing chromosomal packaging events in the distal gut. We validate this method on a range of known phage transduction systems and apply it in a small case study with three faecal samples from healthy human donors. We report detection of dozens of ongoing instances of capsid packaging of chromosomal DNA in key commensal bacteria of the human gut such as *Bacteroides* and *Faecalibacterium*, including evidence of LT and GTA.

## Results

### Discrete peaks of packaged DNA sizes can be observed in the faecal VLP fractions

To test whether discrete peaks of packaged DNA sizes can be recovered from the faecal microbiome, we performed a pilot study on three freshly collected faecal samples from three adult donors (study protocol APC055; donor codes 925, 928, 942). Faecal VLPs were extracted by filtration, concentrated by utracentrifugation and purified using a CsCl density step gradient. The two brightest bands visible in each gradient (further referred to as “B” for bottom, and “T” for top) were collected. After removal of free DNA by DNAse digestion, high molecular weight (HMW) encapsidated DNA was gently extracted to prevent fragmentation. Oxford Nanopore adaptor ligation-based sequencing was used to generate sequencing reads corresponding to the full length of the extracted DNA fragments (**Fig. 1A**). We observed that the length distribution of Nanopore reads included discrete peaks (ranging from ∼4 kb to ∼100 kb) that are likely to correspond to distinct populations of encapsidated DNA originating from different viral agents. The sizes of the DNA fragments in these peaks are consistent with the packaging capacity of small to medium sized tailed phage genomes and were both individual and VLP fraction B or T-specific (**Fig. 1B**). Sample 942T shows signs of possible DNA degradation and enrichment of short fragment sizes. Other samples were dominated by peaks of discrete read lengths with little or no visible DNA degradation.

**Figure 1.**
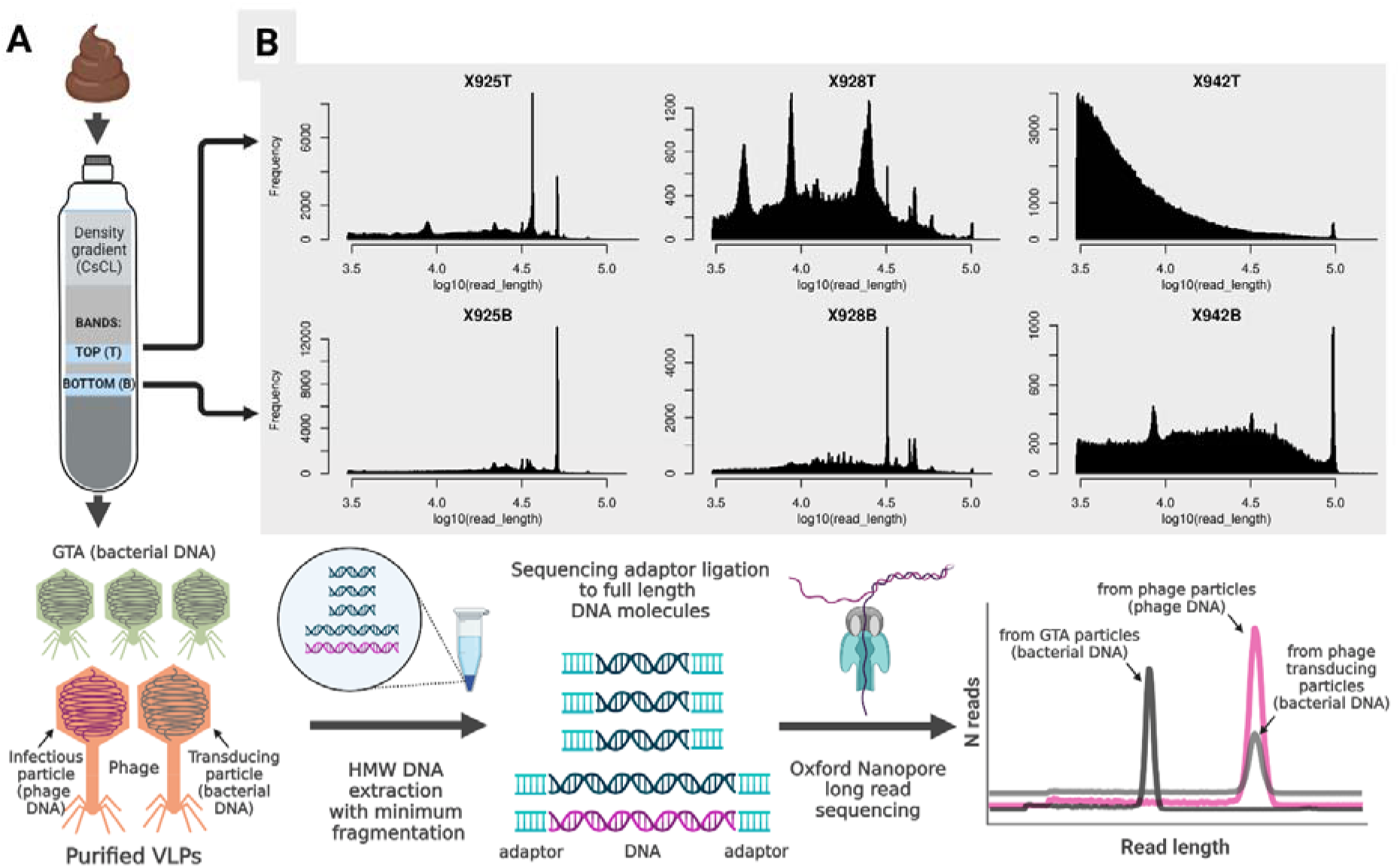
Concept summary of the approach used in the present study. **A,** VLPs are purified from the human faecal samples (n=3), resulting in two VLP fractions (bands) per sample; relatively conserved DNA size packaged by each phage (both phage and bacterial DNA), and GTA leads to narrow peaks of read lengths in the Nanopore full genome-size read sequencing; **B,** distributions of read lengths obtained from each of the six faecal VLP fractions (-B, bottom, heavier particles; -T, top, lighter particles).

To check if our interpretation of the observed pattern of read length peaks is plausible, we applied the same protocol to pure cultures of known transducing/packaging bacterial or bacterium-phage pairs. Sequencing of encapsidated DNA extracted from a *Bacteroides intestinalis* phage crAss001^36^ lysate produced reads of ∼102 kb, aligning end-to-end with the phage genome. No reads aligning to the host chromosome were obtained, suggesting lack of transducing activity in this system (**Fig. S1A**). *Salmonella* phage P22, propagated on *Salmonella enterica* LT2, produced 42.6 kb reads aligning to its 41.7 kbp genome – only slightly shorter than a commonly accepted estimate of the size of packaged DNA of 43.4 kbp^37^. Approximately 4.5% of the total number of long reads (**Fig. S1B**) aligned to dispersed locations in the host chromosome in a pattern consistent with a GT mechanism^12^. The *E. coli* phage P1 capable of GT has a genome of 93.6 kbp and is expected to package 110-115 kbp of terminally redundant DNA ^38^. In our hands, however, 95-96 kb reads were obtained, with 12.8% of them aligning to various locations in the *E. coli* chromosome (**Fig. S1C**). A multi-lysogen strain *Enterococcus faecalis* VE14089 was used as a reference for LT^9,12,39^. VLPs from the *E. faecalis* VE14089 culture contained one induced prophage, pp5, but also contained chromosome fragments of phage genome size adjacent to the prophage location, consistent with the LT model (**Fig. S1D**). Finally, we confirmed that the GTA-like element PBSX in *Bacillus subtilis* packages 13.4 kbp evenly dispersed chromosomal DNA fragments (**Fig. S1E**)^12,40^.

### Taxonomic composition of packaged bacterial DNA in the faecal VLP fractions

Hybrid assembly of short Illumina reads and long Nanopore reads across total DNA and VLP DNA samples yielded a total of 85,015, 42,751 and 45,265 non-redundant metagenomic scaffolds (>1 kb) for donors 925, 928 and 942, respectively. These numbers included 564, 423, and 201 complete and partial viral genomic scaffolds. Based on the results of metagenomic binning, assigned taxonomy (Genome Taxonomy Database, GTDB), and inducible prophage content (detectable in the VLP fraction), all scaffolds were assigned to the following categories: 1) purely viral (mainly bacteriophage) scaffolds (n = 1137 across the three subjects); 2) un-binned bacterial scaffolds (n = 145,909); 3) bacterial scaffolds with prophages (n = 135); 4) bacterial scaffolds binned into 116 metagenomically assembled genomes − MAGs (n = 22,731); 5) MAGs scaffolds with prophages (n = 32); 6) viral scaffolds binned with bacterial MAGs (n = 51). In addition to that, 850 archaeal scaffolds were assembled, with some of them binning into two archaeal MAGs, and none containing identifiable proviruses.

As expected, the sequence space (% of Illumina reads aligned) in the total community DNA was completely dominated by bacterial MAGs and solitary scaffolds, with ≤ 2.7% of reads aligning to purely viral scaffolds (**Fig. 2A**). At the same time, VLP DNA fractions B and T mainly contained sequences mappable to either purely viral scaffolds, or bacterial scaffolds and MAGs containing prophage sequences. Nevertheless, 0.8 – 4.6% of reads in the B fraction and 3.6 – 5.4% of Illumina reads in the T fraction aligned to purely bacterial scaffolds (in or outside of MAGs), indicating either contamination with total community DNA or bacterial DNA packaged into VLPs. However, family level composition of this sub-set of VLP reads differed drastically from that of the total community DNA (**Fig. 2B**). In the total community DNA, families *Bacteroidaceae*, *Rikenellaceae* and *Lachnospiraceae* were the most abundant, whereas in VLP fraction (especially fraction T and donor 942) were dominated by families *Ruminococcaceae* and UBA660 (according to the GTDB taxonomy). To further investigate the origins of this enrichment, we examined the ratios of relative abundance of each of the genomic scaffolds in the VLP fractions versus the total community DNA. The distribution of these ratios deviated from the normal distribution in upper quantiles (p = 1.3e−36, Kolmogorov-Smirnov test), indicating disproportionate enrichment of some of the scaffolds (viral and bacterial) in the VLP fractions (**Fig. S2A**). We arbitrarily set a cut-off of +2 SD to select for enriched scaffolds (**Fig. S2B**) and extracted bacterial MAGs that contained such VLP-enriched scaffolds. As shown in **Fig. S2C**, VLP fraction B mainly contained DNA mapping to individual overrepresented MAG-associated scaffolds containing prophages (genera *Bifidobacterium*, *Dysosmobacter*, *Ruminococcus*). At the same time, VLP fraction T provide examples of enrichment of nearly entire bacterial MAGs belonging to genera *Gemmiger*, CAG-115 (family *Ruminococcaceae*), HGM11616 (family *Christensenellaceae*), and CAG-628 (family UBA660), in line with the composition of dominant bacterial families seen in **Fig. 2B**.

**Figure 2.**
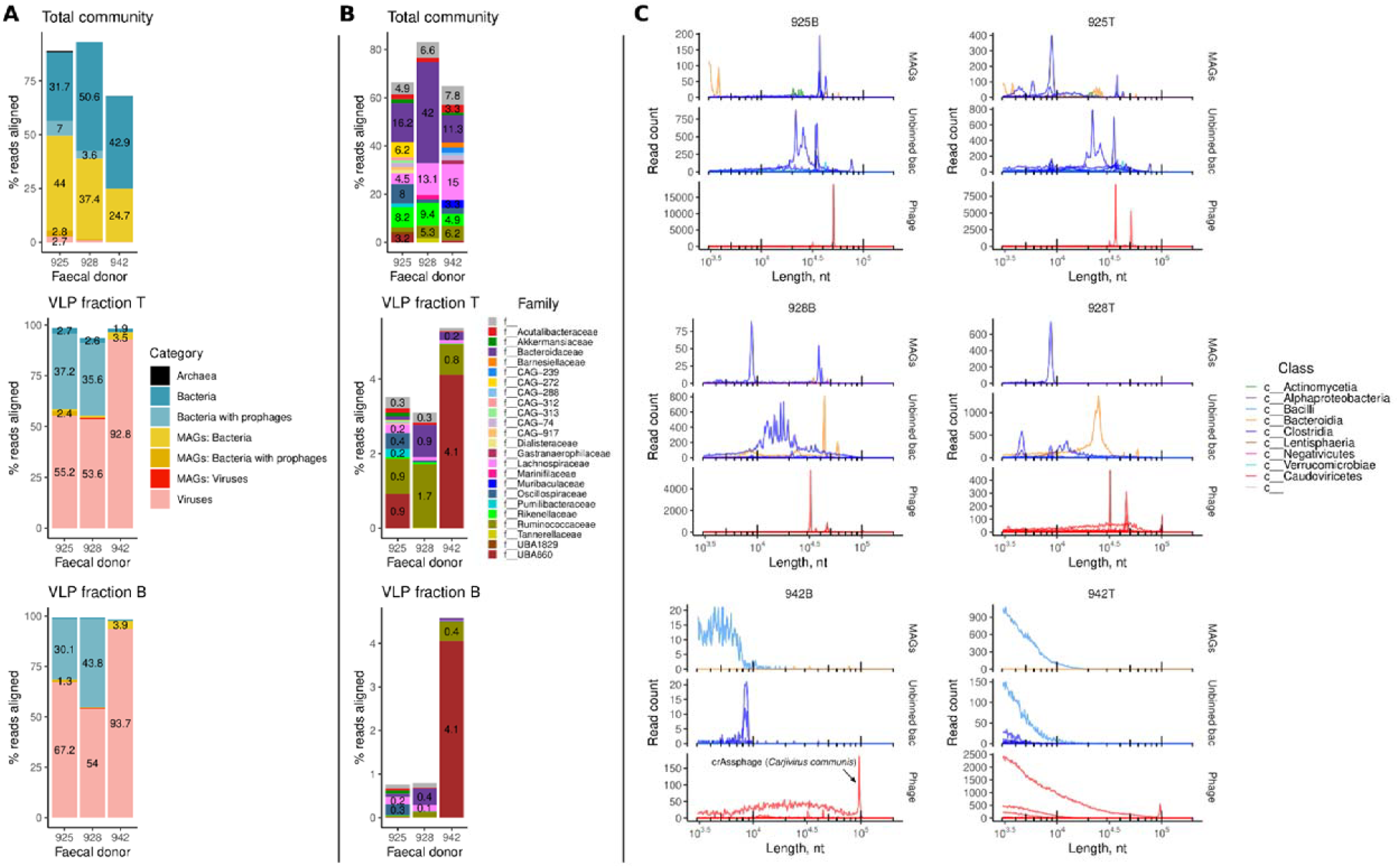
Taxonomic composition of DNA in the faecal VLP fractions and full genome Nanopore read sizes associated with different taxonomic groups. **A**, composition of metagenomic short reads in the total faecal community DNA, as well as two VLP fractions after mapping to metagenomic scaffolds; **B**, family-level composition of bacterial DNA in the same samples; **C**, dissection of Nanopore full genome-size read length distributions by mapping them to various metagenomic entities; each curve is a separate MAG/bacterial species/virus genome; curves are coloured according to taxonomic class (both bacteria and phages) – see text and **Figs. S3-S8** for more details.

### Long-read sequencing of VLP-associated bacterial DNA reveals different transduction modes

We then proceeded to characterise the sequence space of long Nanopore reads obtained from HMW VLP DNA, considering read length distributions. We obtained 0.77 million reads greater than 3kb in length, of which 0.7 million were mapped successfully to genomic scaffolds and assigned to either a bacterial MAG, or, in case of solitary bacterial and viral scaffolds, to the lowest taxonomic rank possible. Reads were then classified into three broad categories: bacterial MAGs (including prophage regions), un-binned bacterial scaffolds (including prophage regions), and viruses. After removal of reads aligning to rare entities, 0.68 million long reads were retained bearing 146 unique taxonomic labels across the three human donors. As shown in **Fig. 2C**, reads associated with specific taxa demonstrated very specific read length distributions, often concentrating into narrow peaks (from 4.6kb to 100kb), with one or more specific narrow peaks per taxa or MAG. Interestingly, peaks representing some of the longest reads obtained in this study (∼100 kb) were mapped to a nearly complete genome of crAssphage (*Carjivirus communis*, 93.1 kb), the most abundant bacterial virus in the human gut^41^.

Manual examination of individual read-length distributions and peaks per each MAG and taxa, as well as examination of read mapping patterns of both the short and long reads to all scaffolds constituting a given MAG or a taxonomic unit in a given sample (**Figs. S3-S8**), allowed us to identify 63 clearly discernible packaging events involving bacterial genomic DNA (including prophages, **Table S1**), across all three donors. Based on read length profiles and mapping to the reference scaffolds, the majority of packaging events associated with the VLP B fraction could be assigned to the induction of prophages (20/31), whereas the remaining cases were consistent with GT, LT and GTA events. The majority (22/32) of events observed in the VLP T fraction were consistent with GTA-like packaging. Some examples of these events are shown in **Fig. 3 and 4**.

**Figure 3.**
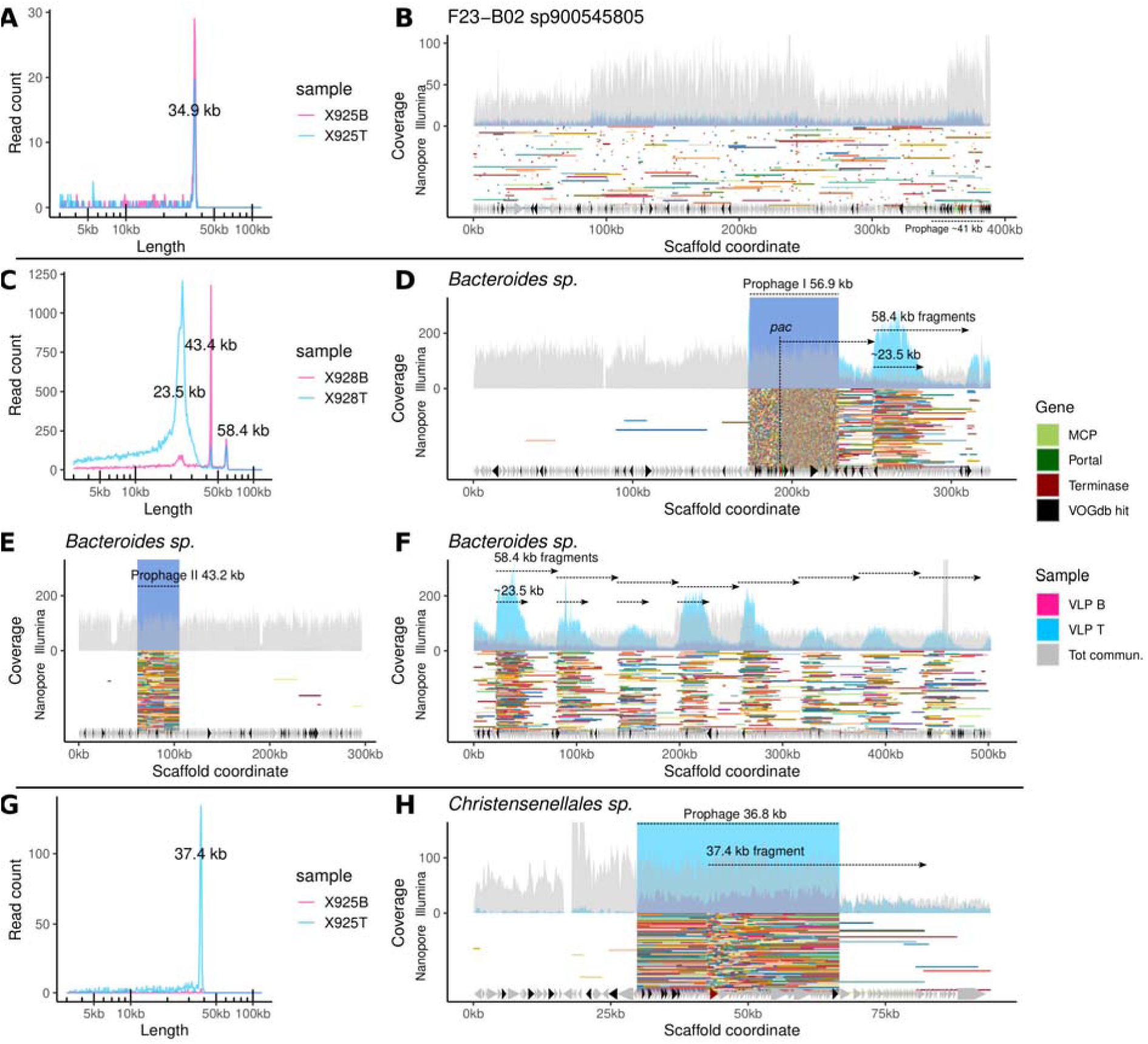
Examples of read sizes and mapping patterns of full-sized Nanopore reads consistent with GT and LT in different gut bacteria. **A** and **B**, distribution of full-sized Nanopore VLP DNA read lengths and their mapping to an uncultured bacterial species’ F23-B02 sp900545805 389 kb genomic scaffold in donor 925 – possible GT; area plots in **B** represent coverage by short Illumina reads, whereas segments of random colours are long Nanopore reads (more scaffold are shown in **Fig. S9**); **C**, three peaks of VLP DNA read lengths associated with a *Bacteroides sp.* bacterium in donor 928; 58.4 kb is a full-sized read length corresponding to a 56.9 kb prophage I and transducing particle DNA (LT); 23.5 kb ones represent partial phage and bacterial DNA fragments (LT) linked with the same prophage I; 43.4 kb reads align to the second predicted 43.2 kb prophage II; **D-F**, mapping of reads from **C** to three metagenomic scaffolds of *Bacteroides sp.* (see text for additional interpretation and **Fig. S10** for more metagenomic scaffolds from *Bacteroides sp.* in donor 928); **G** and **H**, distribution of read sizes and mapping pattern of reads aligning to a 36.8 kb prophage and adjacent chromosomal regions in a *Christensenellales sp.* bacterium in donor 925 (possible LT).

**Figure 4.**
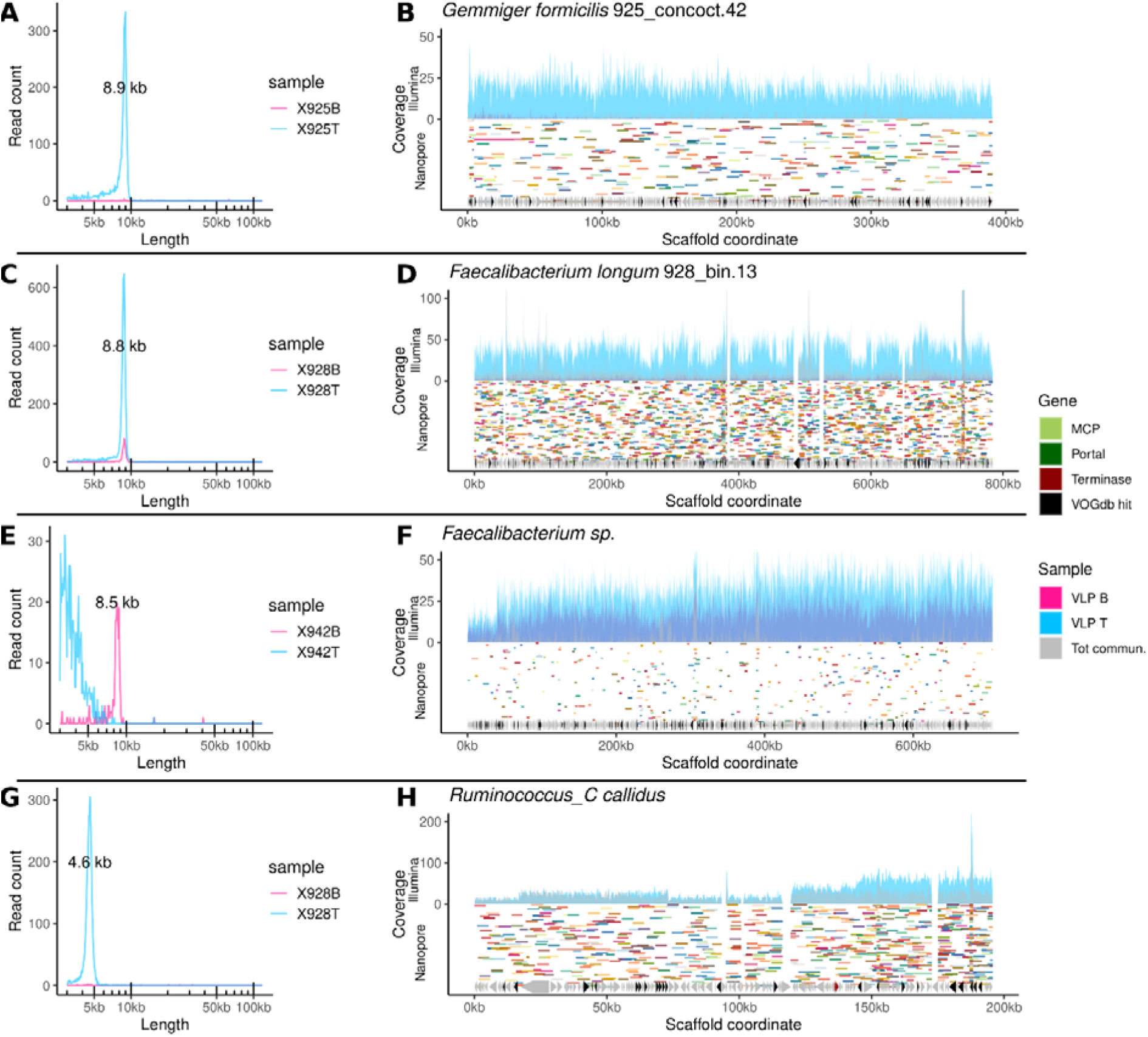
Examples of read sizes and mapping patterns of full-sized Nanopore reads consistent with GTA-like packaging in families *Ruminococcaceae* and *Oscillospiraceae* of gut bacteria. **A** and **B,** distribution of full-sized Nanopore VLP DNA read lengths and their mapping to one of the genomic scaffolds included in MAG 925_concoct.42 (*Gemmiger formicilis*) in donor 925 – possible GTA; area plots in **B** represent coverage by short Illumina reads, segments of random colours are long Nanopore reads; **C** and **D**, same for one of the scaffolds in MAG 928_bin.13 (*Faecalibacterium longum*) from donor 928; **E** and **F**, same for a solitary scaffold from Faecalibacterium sp in donor 942; same for a solitary scaffold of *Ruminococcus callidus* in donor 928.

The first row of panels in **Fig. 3** (**A** and **B**) shows an example of packaging of genomic DNA by a likely GT from an uncultured species F23-B02 sp900545805 (family *Oscillospiraceae*) in the VLP samples from donor 925. Twelve un-binned genomic scaffolds of this species (of which only one is represented in Fig. 3) with a combined length of ∼2.4 Mb (**Fig. S9**) show very uniform coverage by both the Illumina and Nanopore reads, with the latter ones forming a narrow peak of lengths at 34.9 kb, consistent with packaging by phages previously described in this bacterial family, with typical genome sizes of 35-37 kb^42^. The contig shown in **Fig. 3B** in fact contains a prophage of an estimated size of ∼41 kb, however, no evidence of its induction or preferential packaging can be seen, suggesting that a different phage is responsible for the observed packaging. As shown in **Table S1**, additional sets of genomic scaffolds assembled from the same faecal sample were taxonomically annotated as family *Oscillospiraceae* and class *Clostridia*, while showing similar patterns of VLP read lengths as F23-B02 sp900545805, and therefore might in fact originate from the same species. Some of these scaffolds contain complete phage/prophage genomes with genome sizes 33.3-34.5 kb, producing packaged DNA of similar size (34-35kb; **Supplementary Dataset** at DOI 10.6084/m9.figshare.26310658) and can be speculatively implicated in the GT event seen in the species F23-B02 sp900545805.

An example of an LT mechanism is shown in **Fig. 3C-F**. One of the genomic scaffolds belonging to the genus *Bacteroides sp.* in subject 928 contained a highly induced 56.9 kb prophage, producing full-length Nanopore reads (58.4 kb) with random terminal redundancies, as well as shortened reads of ∼23.5 kb, mainly originating from the VLP fraction T (“top” band – less dense phage capsids; **Fig. 3C-D**). As expected, the majority of reads within these two length peaks align to the circularised/concatamerised phage genome. However, a significant fraction of reads within these peaks includes “chimeric” reads, starting from the hypothetical *pac* site located next to the terminase gene and extending into bacterial chromosome outside the prophage region, as well as reads with alignment start sites (multiples of 58.4 kb) located at regular intervals from the prophage *pac* site. For instance, among reads longer than 40 kb, 1,731 aligned within the prophage, 500 aligned outside of the prophage, while 13 reads were chimeric, starting at the putative *pac* site and extending beyond the prophage boundary (**Fig. 3D**). This is consistent with the model of *in situ* prophage over-replication and unidirectional chromosomal DNA packaging that define the LT mechanism. The shorter reads align at regular intervals in such a manner that the alignment start site coincides with the start site for full-length, but the end site is located at random length from the start (∼23.5 kb on average), suggesting that they could have resulted from partial capsid DNA ejection and degradation after packaging, during storage, or during the VLP purification steps. This agrees well with the fact that this type of reads was associated with VLP fraction T (less dense particles).

In the VLP fraction T Illumina short reads, this shortening of packaged DNA fragments creates an effect of periodic waves of coverage, extending from the prophage site. A number of *Bacteroides* genomic scaffolds (n = 10, total length 2.7 Mb), the longest of them being 503 and 669 kb, were recovered from the total community assembly in subject 928 with the same pattern of coverage by Nanopore and Illumina reads (**Fig. 3F**, **Fig. S10**), suggesting very efficient LT-type packaging of a very large portion of the genome, if not the entire genome, in this strain of *Bacteroides sp*. At least one additional inducible 44.2 kb prophage (packaged DNA size 43.4 kb) is likely to be a part of the same genome, but incapable of LT (**Fig. 3E**).

Another example of possible LT, albeit with a lower read coverage, is presented in **Fig. 3G-H**. One out of the three genomic scaffolds from subject 925, belonging to a bacterium in the order *Christensenellales*, contains a 36.8 kb prophage element, with 37.4 kb Nanopore reads aligning to its circular or concatemerized isoform. A few read alignments, however, begin in the vicinity of the terminase gene (possible *pac* site) and extend into the chromosomal DNA downstream of the prophage integration site, while having the same characteristic length of 37.4 kb.

### GTA-like packaging is prevalent in the gut and is primarily associated with families Ruminococcaceae and Oscillospiraceae

The most common mechanism of VLP packaging of bacterial genomic DNA observed in this study was encapsidation of short seemingly random fragments of chromosomal DNA, with relatively even coverage of large spans of genomic scaffolds. This is consistent with the presence of active GTA elements. Examples of this type of DNA packaging were especially observed in the families of gram-positive anaerobic bacteria *Ruminococcaceae* (*Faecalibacterium spp.*, *Gemmiger spp.*, CAG-115) and *Oscillospiraceae* (*Ruminococcus callidus*, *Onthomonas sp.*; **Fig. 4**). In particular, possible GTA-like packaging in *Faecalibacterium spp.* was linked to several MAGs as well as sets of unbinned genomic scaffolds across all three donors, in line with the high relative abundance and prevalence of this microbial genus in the human gut microbiome. The lengths of fragments packaged in *Faecalibacterium spp.*, *Gemmiger spp.* varies over a very narrow range: 8.4-8.9 kb, whereas *R. callidus* provides an example of much shorter fragment packaging of just 4.6 kb.

A search was conducted for GTA-like elements in several MAGs showing patterns of VLP long-read coverage indicative of GTA production (925_bin.12 – *Onthomonas* sp900545815; 925_concoct.42 – *Gemmiger formicilis*; 925_bin.142 – *F. prausnitzii*; 928_bin.13 – *F. prausnitzii*). One of these MAGs, 925_bin.142, contained a strong candidate for a GTA operon. The 10.5 kb operon (GTA I) consists of ten open reading frames, whose predicted products include a possible small terminase subunit (TerS) lacking any obvious DNA binding domain^18^, as well as homologs of a large terminase (TerL), portal protein, head scaffolding and major capsid proteins, as well as several conserved phage-related hypothetical proteins. We conducted a search for homologous regions across all metagenomics scaffolds in this study, as well as in reference genomes of *Faecalibacterium*, *Gemmiger* and *Subdoligranulum* (n = 16). We performed clustering of all predicted protein products using MMseqs2 based on amino acid sequence similarity. Functional annotations of GTA genes from 925_bin.142 were then extrapolated to all members of the corresponding protein clusters (PCs). Regions with concentrations of several putative GTA-related PCs are shown in **Fig. 5A**. Putative GTA operons homologous to the one in 925_bin.142 are present in all twelve *Faecalibacterium* genomes examined (*F. prausnitzii* APC942/30-2, APC918/95b, APC942/8-14-2, APC942/18-1, APC942/32-1, APC923/51-1, APC923/61-1, APC924/119, ATCC 27768, ATCC 27766; *F. hattorii* APC922/41-1; *F. duncaniae* A2-165), but not in *Gemmiger* and *Subdoligranulum*.

**Figure 5.**
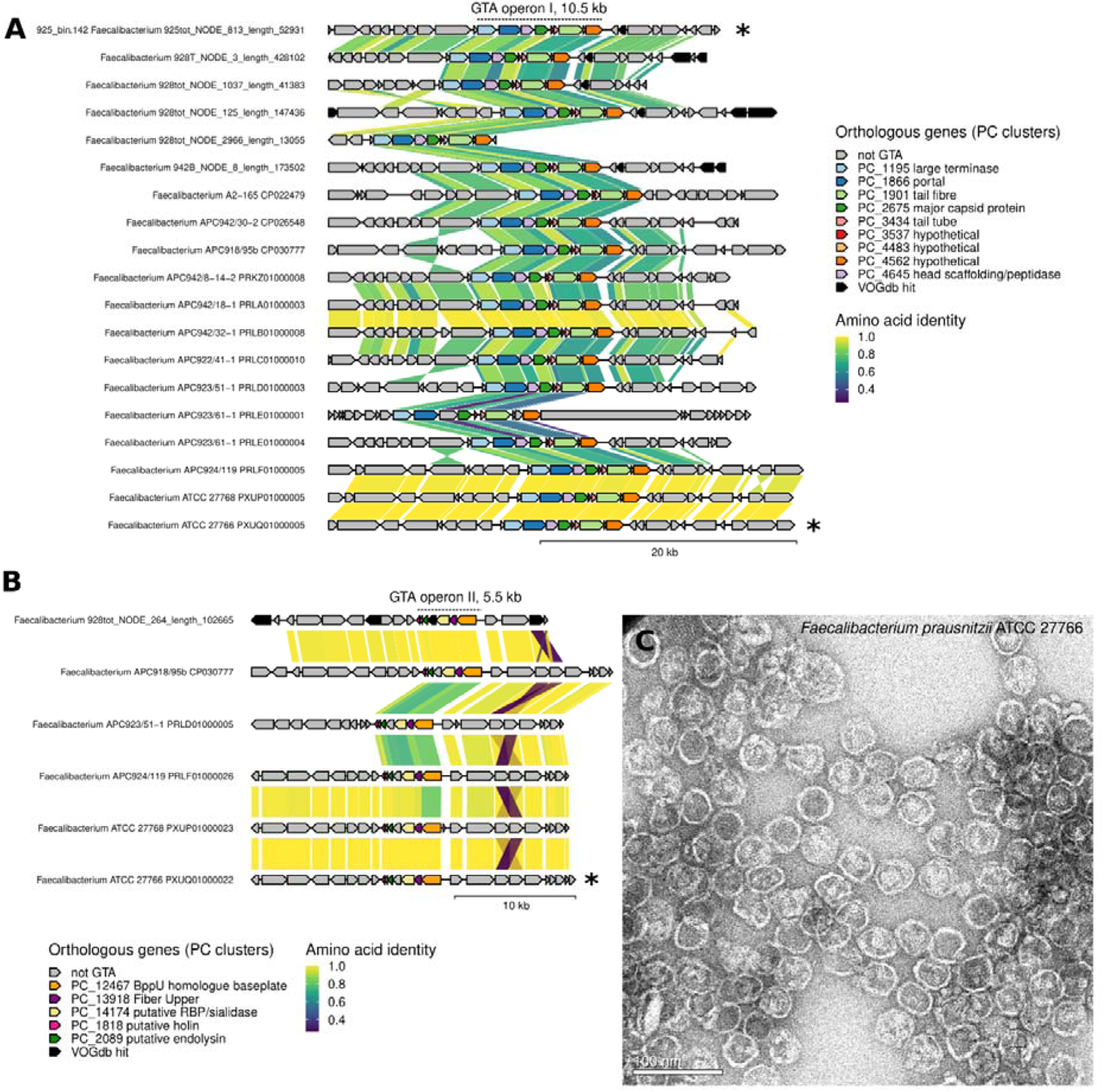
GTA-like elements encoded in MAGs and metagenomic scaffold from *Faecalibacterium spp.* assembled in this study, as well as in genomes from *Faecalibacterium spp.* isolates. **A**, a 10.5 kb candidate GTA operon I was identified of a 52.9 kb metagenomic scaffold included in *Faecalibacterium prausnitzii* MAG 925_bin.142 from donor 925; homologs of this operon are widely distributed in other genomic scaffold from *Faecalibacterium spp.*, as well as in eleven genomes of *F. praunsitzii* isolates (including ATCC 27766), an *F. duncaniae* genome (A2-165) and an *F. hattorii* genome (APC922/41-1); annotated GTA-like structural protein genes are highlighted with colours; other genes with hits to Virus Orthologous Groups Database (VOGdb) are highlighted in black; link colours are based on protein-protein amino acid identity valkues calculated using MMseqs2; **B**, a separate 5.5 kb operon encoding additional structural proteins and lysis genes identified in a *Faecalibacterium sp.* genomic scaffold, assembled in this study, as well as in several *F. prausnitzii* isolates, including ATCC 27766; **C**, Trasmission electron microscopy of capsid-like structures (∼40 nm in diameter) spontaneously produced in broth culture of *F. prausnitzii* ATCC 27766.

Anaerobic broth cultures of some of these strains (APC923/51-1, ATCC 27768, ATCC 27766) were prone to lyse in the late stationary phase (48h-72h). A culture supernatant collected from late stationary phase *F. prausnitzii* ATCC 27766 was concentrated and separated using CsCl step gradient ultracentrifugation, producing a single band. Negative stain TEM imaging revealed that the band contained capsid-like particles with an approximate diameter of 40 nm (**Fig. 5C**), which is consistent with a potential T=3 capsid architecture and packaging capacity of 8.5 kb DNA if the DNA is packaged in equivalent density as in Rc and CcGTA^26,35^. The capsid particles have no clearly identifiable tails, but the tails may have been lost during purification or are very short. Indeed, the GTA I cluster encodes proteins homologous to those used by short-tailed phages. LC-MS proteomics analysis of these particles demonstrated that structural proteins encoded by the GTA I operon (e.g. major capsid protein RCH50849.1, and portal protein RCH50851.1) constitute the majority of all ATCC 27766 proteins present in the sample (**Table S2**). Moreover, a number of phage-like structural proteins encoded in the ATCC 27766 genome outside of GTA I operon were detected. When mapped to the genome of ATCC 27766, these formed a second operon (GTA II, 5.5 kb), comprising structural protein genes: baseplate, tail fibre, putative receptor-binding protein (RBP), as well as the lysis proteins – holin and endolysin. No other putative prophage proteins were detected that could account for the encapsidated DNA. Taken together, these two operons probably constitute most, if not all of the functions needed for GTA particle morphogenesis^17^ (most likely podovirus-like), DNA packaging and cell lysis in *F. prausnitzii* ATCC 27766. GTAs often require auxiliary structural or maturation genes, and we cannot exclude that there are more to be identified here. Both GTA loci are also present in strains APC924/119, APC923/51-1, APC918/95b, ATCC 27768, but GTA locus II was not detected in the remaining seven strains. It is likely that the strains that only carry GTA operon I, have either lost GTA operon II or possess a more divergent set of genes that were overlooked by our analysis. In the strain APC918/95b, for which a complete circular genome sequence is available (CP030777), the two operons (GTA I: 2536737-2546346 and GTA II: 2672270-2677176) are separated by a distance of ∼125 kb.

## Discussion

We employed a long-read single molecule nanopore sequencing-based approach to identify capsid-mediated packaging and mobilisation of bacterial DNA in the human gut microbiome. The size of the packaged single DNA molecule adds an important extra layer of information that allows us to distinguish between encapsidated DNA (producing narrow peaks of read lengths, indicative of the size and type of capsids) and contaminating and degraded bacterial DNA. This approach also allows individual capsid-packaged DNA molecules to be traced, thus allowing us to determine their origin, chromosomal location, structure of ends, to identify circularly permutated (headful packaging) and chimeric reads (ST and LT), and in some cases to link transducing particles with the likely origin phages. This approach is also uniquely suitable for identifying instances of host DNA encapsidation involving GTAs and GTA-like elements. GTAs can be challenging to study because the GTA-encoding genes are not selectively packaged but rather the entire host genome is packaged relatively uniformly^12^.

Our results from three human faecal samples show extensive packaging of host DNA, representing up to 5.4% of the total VLP DNA content. This was broadly representative of all main bacterial phyla present in the human gut, and included examples of possible GT, ST, LT, and GTA. Ongoing packaging events involving phyla *Bacillota* (GT, GTA) and *Bacteroidota* (LT) were especially prevalent. This agrees with numerous reports providing evidence of a high rate of HGT, including phage-mediated HGT in the human gut^6,7^, and its central role in the evolution of complex microbial communities on ecological time scales^43,44^. Interestingly, a recent study examining the rates of inter-species HGT in the human gut demonstrated that class *Clostridia* (phylum *Bacillota*) and order *Bacteroidales* (phylum *Bacteroidota*) show the highest rates of horizontal gene transfer^45^. An earlier study demonstrated extensive HGT between *Bacteroidales* species in the human gut involving integrative and conjugative elements (ICE)^46^.

The majority of bacterial packaging events in our dataset were prophage induction events. It has been previously shown that lysogens are prevalent in the gut bacteriome^47,48^ and that temperate phages dominate in the gut virome^49,50^. Furthermore, the gut environment is conducive to prophage induction^51,52^ through signals produced by the host organism, resident bacteria, and external sources (antimicrobial peptides, quorum sensing autoinducers, immune and hormonal signalling, nutrients, bile salts, etc.)^53^, which subsequently affects transduction by temperate phages and, likely, GTAs that are also inducible through the SOS response mechanism^17^.

One of the interesting finds in our study was a clear signature of prophage-driven LT in *Bacteroides*, the most abundant genus of bacteria in the human gut microbiome^54^. LT was initially discovered in temperate phages of *S. aureus*^9^ and subsequently in *S. aureus* PICI^11^. It was later observed in *E. faecalis*^12^ and *Salmonella*^13,14^, showing that lateral transduction is neither unique to *Staphylococcus*, nor to the phylum *Bacillota*. In this study we expand the taxonomic range of LT-capable phages even further, with examples of lateral transduction observed in *Bacteroidales* (phylum *Bacteroidota*) and *Christensenellales* (phylum *Bacillota*).

While it is evident that phage involvement in genetic exchange can provide a benefit to the bacterial population^44,55,56^, it appears to be costly to the phages in that it decreases the number of infectious particles. This raises the questions of why bacteriophages often engage in host DNA mobilisation, seemingly to the detriment of their own spread in the population. One study suggests that GT confers fitness advantage to temperate phages in rapidly changing environments and can be seen as an adaptation inherent to their life cycle^57^. Transduction by temperate phages can promote adaptations and increase survival of the lysogenized subpopulation while eliminating non-lysogenized competition, thus ensuring vertical transfer of the prophage^58^. A recent study shows that the ability to package host DNA in *Salmonella pac*-type phages evolved distinct generalised transduction strategies^14^, similarly indicating that host DNA mobilisation is a consistently conserved adaptation rather than a persistent mistake.

The most striking observation was evidence of GTA operating at high efficiency in members of the families *Ruminococcaceaa* and *Oscillospiraceae*, including genus *Faecalibacterium* – another highly abundant microbial genus in the human gut. Very few GTA elements have been described so far^17,35,59–61^. GTA production has been observed in the human pathogen *Bartonella bacilliformis*^62^, and GTA-like gene clusters were found in *Brucella abortus* and *Brucella suis* genomes^60^, but not in human gut commensals. Our understanding of GTA operation, prevalence and phylogeny remains extremely limited, with formal recognition and taxonomic classification of GTAs only recently accepted, and encompassing primarily GTAs in Alphaproteobacteria and Spirochaetia (*Brachyspira*)^15,16^. The few known GTAs outside of Alphaproteobacteria have been identified based on their random DNA content. The GTA loci typically remain either unknown or incomplete and thus no taxonomic classification can be attempted^23,24,63^.

GTA production could potentially play a role in genetic exchange in *Faecalibacterium*, where GTA operons appear to be near-ubiquitous. Our results show that at least some of these operons are active and produce GTA particles, but further research is needed to determine whether the particles are capable of successfully delivering their transducing DNA, or if they instead function similarly to the GTA-like element PBSX in *Bacillus subtilis*^40,64^, which does not inject packaged DNA but acts as a tailocin^65^. If *Faecalibacterium* GTAs are indeed a functional transduction system, they could confer fitness advantage beyond adaptive trait acquisition. It has been shown that in *Caulobacter crescentus* gene transfer by GTAs improves cell survival in stationary phase and after DNA damage by enabling DNA repair through homologous recombination^26^. A previous comparative analysis of *Faecalibacterium* genomes demonstrated strikingly low synteny between available genomes, indicating high genome plasticity, as well as high prevalence of ICE^66,67^. Whether or not the frequent genetic exchange between individual strains via GTA contributes to high genome plasticity and low genomic synteny in the genus *Faecalibacterium*, with or without the involvement of ICE, remains to be established. However, it seems logical to propose that in a complex and dense microbial community, such as the human gut microbiome, various modes of capsid-mediated DNA mobilisation (GT, LT, GTA) can interact with other HGT mechanisms (transposons, integrons, ICE, plasmids) as parts of an integrated and highly active network of within- and between-species genetic exchange.

### Limitations of the study

The approach we present relies on ligation-based long-read sequencing of native DNA molecules and requires large amounts of high quality, high purity, high molecular weight DNA. The method of DNA extraction and library preparation is challenging and requires considerable amounts of freshly collected starting faecal material, limiting application of the method in large scale population studies. In addition, these factors limit the depth of long read sequencing meaning that some coverage patterns and size distributions can be inconclusive, and some packaging events may be missed entirely.

Additionally, extracellular vesicles (EVs) can potentially be co-purified with capsids. In particular, this might create some difficulties in discrimination between EV and GTA, as gene transfer agents exclusively and non-selectively package bacterial DNA resulting in read coverage patterns similar to those of EV DNA. While EVs are an emerging field in gut microbiome studies^68^, considerable research has been done into bacterial EVs in marine ecosystems. It has been suggested that in marine viromes extracellular vesicles, rather than GTAs, represent the main source of packaged non-viral DNA^30^. Broader DNA size distribution typical for extracellular vesicles^69^ makes it possible to distinguish between GTA and EV DNA using size analysis. However, for example, *Prochlorococcus* and *Streptomyces* EVs show presence of discrete DNA ‘bands’ of certain size^70,71^.

## Acknowledgements

The authors thank Maria Chechik (University of York) for performing VLP imaging using TEM and the York-JEOL Nanocentre at the University of York for the instrument access and technical support; and York Centre of Excellence in Mass Spectrometry for conducting mass spectrometry analysis (supported by a major capital investment through Science City York and Yorkshire Forward with funds from the Northern Way Initiative and EPSRC [EP/K039660/1; EP/M028127/1]). The authors thank Prof. Breck A. Duerkop (University of Colorado) for providing some of the bacterial and phage strains used in this study.

This research was funded by the European Research Council (ERC), under the European Union’s Horizon 2020 research and innovation programme (grant agreement No. 101001684 – PHAGENET). Andrey Shkoporov and Pavol Bardy were funded by Wellcome Trust Research Career Development Fellowship [220646/Z/20/Z] (A.S.) and Sir Henry Wellcome Postdoctoral Fellowship [224067/Z/21/Z] (P.B.). Jason Wilson is funded by a BBSRC responsive mode grant [BB/V016288/1] and Paul Fogg is jointly funded by a Wellcome Trust and Royal Society Sir Henry Dale Fellowship [109363/Z/15/A] and a BBSRC responsive mode grant [BB/V016288/1].

This research was funded in whole, or in part, by the Wellcome Trust [220646/Z/20/Z, 224067/Z/21/Z and 109363/Z/15/A]. For the purpose of open access, the authors have applied a CC BY public copyright licence to any Author Accepted Manuscript version arising from this submission.

## Author contributions

Conceptualization, T.B. and A.N.S.; Methodology, T.B. and A.N.S.; Investigation, T.B., J.S.W, P.B., M.S., Conor H., E.V.K.; Writing – Original Draft, T.B. and A.N.S.; Writing – Review & Editing, T.B., J.S.W, P.B., B.G., P.C.M.F., C.H. and A.N.S.; Funding Acquisition, A.N.S.; Resources, B.G. and A.N.S.; Supervision, P.C.M.F., C.H. and A.N.S.

## Data availability statement

Raw sequencing data is deposited at NCBI SRA database under a BioProject accession PRJNA1135972. Supplementary dataset, which includes MAG and genomic scaffold sequences, read mapping data, a complete set of R scripts and input data needed to recreate all tables and figures in the manuscript, and a complete set of raw figures describing read length distributions and genomic scaffold mapping patterns of long Nanopore reads is available at FigShare under DOI 10.6084/m9.figshare.26310658.

## Materials and Methods

### Preparation of VLPs from human faecal samples

Ten grams from each of the faecal samples were subdivided into 4 aliquots of 2.5 g each. 20 ml of SM buffer were added to each aliquot and homogenisation was achieved by vortexing for 5 min. Each tube was topped up with SM buffer to a final volume of 45 ml and vortexed for and additional minute. The tubes were then centrifuged at 10,000 rpm for 20 min at 4°C. The supernatant was decanted and centrifuged a second time as above. The supernatant was passed through a series of syringe mounted PES Filtropur filters; once through 0.8 µm pore membranes and twice through 0.45 µm pore membranes. The filtered supernatant was decanted into 10 mL polycarbonate ultracentrifuge bottles (Thermo Scientific) and centrifuged at 40,000 rpm at 4°C for 2 hours, in a F65L-6x13.5 rotor in Sorvall™ WX+ ultracentrifuge (Thermo Scientific). The supernatant was discarded and the pellets from each tube was resuspended in a total of 5 ml of SM buffer, followed by additional filtration through a 0.45 µm pore syringe filter. Five ml of the supernatant were then layered over a CsCl step gradient (4 ml 3M and 4 ml 5M CsCl solution in SM buffer) in 13.5 mL Quick-Seal Ultracentrifugation Tubes (Beckman Coulter). Tubes were sealed and centrifuged in an F65L-6x13.5 rotor at 37,000 rpm at 4°C for 2.5 hours. The VLP material produced multiple bands with two major bands visible in all three faecal samples, which were collected using a hypodermal needle and de-salted by three rounds of concentration-dilution with SM buffer (50mM Tris-HCl pH7.5, 100mM NaCl, 8.5mM MgSO4) on a 100 kDa MWCO Vivaspin (Sigma-Aldrich) centrifugal filter pre-equilibrated with SM buffer, each round at 4,000 g at room temp, followed by concentrating to a final volume of ∼400 µl.

### Preparation of VLPs from model phage-host pairs

*Bacillus subtilis* GTA-like agent PBSX particles were obtained as described in Kleiner et al., 2020. The culture was incubated at 37°C until OD600 reached 0.5. Mitomycin was then added (0.5 µg/ml final concentration) to induce prophage excision, and the culture was incubated for a further 10 minutes, before being washed and resuspended in fresh culture medium. The pellet was thus washed 2 times, and the culture was incubated for an additional 4h. The culture was then centrifuged, and the supernatant recovered and filtered through 0.45 µm PES syringe filters (Filtropur).

P22 and P1: as described in Kleiner et al., 2020^12^; P22 and P1 were propagated overnight in 0.4% LB agar overlay on hosts *Salmonella enterica LT2* and *Escherichia coli MG1655*, respectively. Top agar was collected into 3 ml of SM buffer per plate, vortexed and mixed on a rotary mixer for 30 minutes, centrifuged at 4°C at 3000 g. The supernatant was then filtered through a 0.45 µm syringe filters.

crAss001: propagated overnight in FAB (Neogen) 0.4% agar overlay on host *Bacteroides intestinalis* APC919/174. Filtrate was obtained from top agar as described above.

*E. faecalis* prophage*s*: as described in Kleiner et al., 2020^12^. *Enterococcus faecalis VPE14089* was inoculated at 1% from an overnight culture, grown to an OD600 of 0.5, induced using ciprofloxacin at a final concentration of 2 µg/ml and grown for a further 4 hours. The culture was then centrifuged at 4700 rpm and the supernatant filtered through a 0.45 µm syringe filters.

*Faecalibacterium prausnitzii*: 72h culture of *F. prausnitzii ATCC27766* was centrifuged at 4700 rpm at 4°C for 30 minutes, the pellet was discarded, and the supernatant centrifuged for another 30 minutes. The resulting supernatant was filtered through a 0.45µm syringe filter. NaCl / PEG8000 concentration and chloroform treatment were performed as described above. The resulting aqueous phase was further purified and concentrated to 5 ml by running 2x volumes of SM buffer through a 15ml 100 kDa Amicon column (centrifugation at 3000 g at 18 °C).

### Purifications of VLPs

Filtrates of all cultures were concentrated using 10% w/v PEG 8000 in 0.5M NaCl, precipitated overnight at 4°C and pelleted at 4700 rpm for 20 minutes at 4°C. The pellet was resuspended in 5 ml of SM buffer and equal volume of chloroform was added, mixed by vortexing, and centrifuged at 3000 g for 10 minutes at 18 °C. The aqueous phase was extracted, and chloroform treatment repeated a second time. VLP purification by CsCl gradient centrifugation was performed as described above. A single visible band was collected for DNA extraction.

### High molecular weight VLP DNA extraction

40 ul of 10X DNAse buffer (100mM MgCl_2_, 500mM CaCl_2_), 9 µl (2 U/µl) TURBO™-DNase, and 2 µl (1mg/mL) RNase A (Invitrogen™) were added to each 400 µl sample to remove non-encapsidate DNA and RNA. After incubation at 37°C for 1h, endonucleases were inactivated at 70°C for 10 minutes. Two microlitres of proteinase K (20 mg/ml) and 20 µl of 10% SDS were added, and the sample was incubated for a further 20 minutes at 56°C, followed by the addition of 100 µl phage lysis buffer (4.5 M guanidinium isothiocyanate, 44 mM sodium citrate, 0.88% sarkosyl and 0.72% 2-mercaptoethanol, pH 7.0) and incubation for another 10 minutes at 56°C. An equal volume of phenol:chloroform:isoamyl alcohol (25:24:1) was added and mixed by manually inverting for 2 minutes. Following centrifugation at 8000 g at room temperature, aqueous phase was transferred into a sterile tube. The aqueous phase was gently collected using wide-bore pipette tips, mixed with an equal volume of chloroform, and centrifuged at 8000 g at room temperature for 10 minutes. Sodium acetate (to a final concentration of 0.3M) and 1 µl of glycogen (20 mg/ml) were added to the collected aqueous phase. An equal volume of pre-chilled (–20°C) isopropanol was added and, after gentle mixing, the sample was incubated at –20° C for 30 minutes. Following centrifugation at 17000g at -9°C for 30 minutes, the supernatant was discarded, and the pellet was briefly washed with 1 ml of 80% ethanol. The ethanol was removed, and the pellet was left to dry, prior to being resuspended in 50 μl of nuclease-free water overnight at 4°C. DNA concentration was measured with dsDNA Broad Range Qubit (Invitrogen).

### Total faecal DNA extraction

Total DNA was extracted from the faecal samples as described in Shkoporov et al., 2019. Briefly, 5 g of faecal sample were resuspended in 25 mL of InhibitEx Buffer (Qiagen) and aliquoted into 2 mL tubes containing a mixture of homogenization beads (Thistle Scientific/Biospec Products): 200 μl 0.1 mm zirconium beads, 200 μl 1 mm zirconium beads, one 3.5 mm glass bead per tube. Aliquots were homogenized in the FastPrep-24 Classic bead beater (MP Biomedicals) for 30 seconds, chilled on ice for 1 minute, followed by another round of homogenization. The samples were incubated for 5 minutes at 95°C, followed by DNA extraction using QIAamp Fast DNA Stool Mini kit according to the protocol. DNA was quantified using Qubit dsDNA BR kit.

Selection of large (>3Kb) DNA fragments was achieved by extraction from excised agarose gel band following electrophoresis. The DNA was extracted from the gel band by running electrophoresis with the band enclosed in a 14kDa dialysis tubing. The DNA suspended in TAE buffer was precipitated with isopropanol as described above.

### DNA library preparation and sequencing

DNA quality was assessed using Nanodrop 1000 spectrophotometer. DNA was considered of acceptable quality if 230/280nm and 260/280nm absorbance ratios were ∼1.8 and ∼2-2.2 respectively. Sequencing libraries were prepared using Nanopore Ligation sequencing kit LSK109. Sequencing was performed on R 9.1.4 MinION flowcell (FLO-MIN114).

For short read sequencing, 10-100 ng DNA in 50 ul deionised water was sheared using Covaris M220 focused ultrasonicator with the following parameters: time 35 seconds, peak power 50.0, duty factor 20.0, cycles/burst 200. Fragmented DNA was concentrated to 15 ul using Genomic DNA clean and concentrator kit (Zymo Research). Sequencing libraries were prepared using Accel-NGS 1S Plus DNA Library Kit (Swift Biosciences). Sequencing was performed with 2x150bp paired end configuration on Illumina NovaSeq 6000. Total number of reads after quality filtration was 81-101 million per VLP DNA sample and 8-71 million per total faecal DNA sample.

### Nanopore reads from model phage-host pairs

Nanopore reads were basecalled using ont-guppy v6.4.6 (in MinKNOW GUI v22.12.7). FASTQ files were converted into FASTA using seqret in EMBOSS v6.6.0.0 and BLASTn ((v.2.10.0+; *- evalue 1e-20*)) searched against the reference genomes of bacteriophages and their bacterial hosts. NCBI RefSeq accession numbers are as follows: *Bacteroides intestinalis* APC919/174 substr 8W, NZ_CP064940.1; Bacteroides phage crAss001, NC_049977.1; *Escherichia coli* str. K-12 substr. MG1655, NC_000913.3; Enterobacteria phage P1, NC_005856.1; *Salmonella enterica* subsp. *enterica* serovar Typhimurium str. LT2, NC_003197.2; *Salmonella* phage P22, NC_002371.2; *Enterococcus faecalis* strain VE14089, NZ_CP039296.1; *Bacillus subtilis* subsp. *subtilis* ATCC 6051, NZ_CP034484.1. Only BLASTn alignments of ≥ 1kb were kept. In case of ambigous alignment of Nanopore reads to more than one genomic scaffold, a scaffold producing the greatest sum of alignment lengths with a given read was considered as correct template. Read length was determined using infoseq in EMBOSS v6.6.0.0.

### Assembly of faecal VLP DNA shotgun reads

Nanopore reads were basecalled using ont-guppy v3.4.4. No further filtration or trimming was applied. Illumina reads (Accel-NGS 1S libraries) were trimmed with cutadapt v2.8 and further trimmed and filtered using TrimmomaticPE v0.39 to remove low quality bases and reads that ended up being too short (*SLIDINGWINDOW:4:20 MINLEN:60 HEADCROP:10*). Hybrid assemblies of filtered Illumina reads and raw Nanopore reads were out carried separately for each of the samples (top and bottom VLP fractions, three donors, n = 6) using SPAdes v3.13.1 in --*meta* mode.

### Assembly of total faecal DNA shotgun reads

Nanopore reads were generated and basecalled using ont-guppy v6.4.6 (in MinKNOW GUI v22.12.7) using default settings for the flowcell and chemistry used. Illumina reads from Accel-NGS 1S libraries were filtered and trimmed using fastp v0.20.0 with the following parameters: *-- length_required 60 --detect_adapter_for_pe --trim_tail1 2 --trim_tail2 2 --trim_front1 8 -- trim_front2 13*. To remove human host contamination reads were subjected to a Kraken2 (v2.0.8- beta) against a custom in-house database consisting of human, mouse, pig and rhesus macaque genomes (Hsap_Mmul_Sscr_Mmus_custom_db). Unclassified reads were retained for assembly. Illumina reads originating from Nextera XT libraries were filtered and trimmed using TrimmomaticPE (*SLIDINGWINDOW:4:20 MINLEN:60 HEADCROP:10*) followed by a Kraken2 search against Hsap_Mmul_Sscr_Mmus_custom_db. Hybrid assemblies of filtered Illumina reads (Accel-NGS 1S library + NexteraXT library) and raw Nanopore reads were out carried separately for each of the three faecal donors using SPAdes v3.13.1.

### Scaffold binning and selection of microbial MAGs

Scaffolds assembled from both the total community DNA and the VLP DNA (two fractions) were pooled together by faecal donor. Scaffolds were filtered to remove sequences less than 1kb long and made non-redundant after an all-vs-all BLASTn (v.2.10.0+; *-evalue 1e-20*) by removing duplicates (shorter scaffolds having 90% of their length represented within longer scaffolds at 99% sequence identity). This procedure yielded 85,015; 42,751; 45,265 non-redundant scaffolds (microbial and viral) for donors 925, 928 and 942. The combined length of metagenomic assemblies were 404, 254 and 158 Mb respectively. Total community metagenomic Illumina reads (Accel-NGS 1S libraries) were mapped back to these assemblies, to determine individual scaffold coverage, using Bowtie2 (v2.3.5.1; *--end-to-end* mode), followed by converting SAM-format alignment files into BAM files, sorting and indexing alignments with Samtools v1.10. Scaffold coverage data was summarised using the program “jgi_summarize_bam_contig_depths” from the MetaBAT2 package. Several suits of tools (MetaBAT2 dev build 2023-03-17, MaxBin2 v2.27; CONCOCT v1.1.0) were applied to bin scaffolds into candidate MAGs with DAS Tool (v1.1.6) being used to integrate binning results and create a non-redundant set of bins (MAGs) for each of the three faecal donors. MAGs were quality checked using CheckM v1.2.2 and taxonomically classified with GTDB-Tk v2.3.2 (workflow *classify_wf*) using GTDB database release 214. MAGs showing completeness of ≤ 50 and contamination of ≥ 5 were discarded, resulting in 70, 33 and 15 high-quality MAGs from donors 925, 928 and 942, respectively. Additionally, taxonomic annotation was performed on all non-redundant genomic scaffolds, regardless of binning result using MMseqs2 *easy-taxonomy* workflow and GTDB database (v214.1) indexed for MMseqs2 search. Functional annotation of all non-redundant scaffolds was performed using DRAM (v1.4.6, database version as of 2023-03-23)

### Selection of viral genomic scaffolds

Viral genomes/genome fragments were identified using two passes of VirSorter2 v2.2.4 on scaffolds assembled from top and bottom VLP fractions. Following the first pass (*--keep-original-seq --min-length 3000*), candidate viral scaffolds completeness and contamination (provirus state) were assessed using CheckV v1.0.1 in the *end_to_end* mode. Provirus and virus sequences identified using CheckV were combined and subjected to a second pass of VirSorter2 with the following parameters: *--seqname-suffix-off --viral-gene-enrich-off --provirus-off --prep-for-dramv -- min-length 3000*. The following additional filtering criteria were applied to VirSorter2 selected scaffolds to qualify as viruses: viral genes count > 0; host genes absent; max_score > 0.95; number of hallmark genes > 2. Additionally geNomad v 1.6.1 (*end-to-end* mode) was employed to identify viruses and plasmids. The combination of two tools allowed to identify 846, 724 and 314 complete and partial viral genomic contigs in donors 925, 928 and 942, respectively. Taxonomic classification of viruses was based on geNomad taxonomic assignments (broad level) and ICTV species-level taxonomy where possible, obtained by BLASTn (*-evalue 1e-20*) search against ICTV exemplar genomes listed in the ICTV VMR resource as of 2022-10-13. Hits were considered the same species if 85% of the scaffold of interest (combined length of BLASTn HSPs’s) were contained within a reference genome at 95% sequence identity level. Additionally, virus scaffolds were searched against total community scaffolds to identify cases where virus scaffolds (one or multiple fragments, e.g. prophages) were contained within larger total community scaffolds (BLASTn, with a threshold at 99% identity and 90% of total length aligned).

### Mapping Nanopore reads to non-redundant scaffolds

Nanopore reads of VLP DNA (top and bottom VLP fractions separately) were mapped to non-redundant scaffolds, separately, in each faecal donor, using Minimap2 v2.17-r941 and *-ax map-ont* preset. SAM alignment were converted into BAM files, sorted and indexed using Samtools v1.10. The *bamtobed* command from bedtools v2.27.1 was used created BED files with Nanopore read alignment coordinates.

### Identification of putative GTA genes within F. prausnitzii 925_bin.142 and their homologues

The 925_bin.142 *F. prausnitzii* MAG sequence was submitted to geNomad^72^ server (https://portal.nersc.gov/genomad/), Phaster server (https://phaster.ca/)^73,74^, and Virsorter2^75^ with a minimum length of 1500. Automatically identified phage regions were manually inspected for the presence of putative GTA gene features. A large terminase (CDS16 in the 52.9 kb genomic scaffold “NODE_813_length_52931_cov_10.520009”) was identified by all three annotation programs, and an upstream putative small terminase (CDS15) that was missed by the programs was identified by manual inspection. CDS15 was submitted to Collabfold v1.5.5.5 structural prediction, with no template and 3 recycles. Eight additional phage-like genes were highlighted in the same 10.5 kb region (GTA I operon) by the annotation software. These included a portal, capsid and some tail components also identified in the MS data. As expected, no DNA replication or phage metabolism genes were identified. To identify possible homologs of these genes in related genome, a total of 16 complete and partial reference genomes of genera *Faecalibacterium*, *Gemmiger* and *Subdoligranulum* were included: ten *F. prausnitzii*, APC942/30-2 CP026548, APC918/95b CP030777, APC942/8-14-2 PRKZ0100, APC942/18-1 PRLA0100, APC942/32-1 PRLB0100, APC923/51-1 PRLD0100, APC923/61-1 PRLE0100, APC924/119 PRLF0100, ATCC 27768 PXUP0100, ATCC 27766 PXUQ0100; *F. duncaniae* A2-165 CP022479; *F. hattorii* APC922/41-1 PRLC0100; *G. formicilis* ATCC 27749 FUYF0100; *G. gallinarum* DSM 109015 JADCKC01; *Subdoligranulum variabile* DSM 15176 CP102293; and *Subdoligranulum sp.* APC924/74 PSQF0100. Translations of all coding sequences, together with translations of all coding sequences in non-redundant scaffolds from the three donors annotated by DRAM, were pooled and subjected to clustering using MMseqs2 (*cluster* command, default settings).

### EM and proteomics

For negative stain electron microscopy, four µl of the purified VLP sample were applied to glow-discharged Formvar/carbon 200 mesh copper grids (Agar Scientific) and stained using 2 % uranyl acetate. After complete drying, the grids were imaged using JEOL 2010 TEM operated at 200 kV at York JEOL Nanocentre.

For MS analysis, 100 µL of the purified material was taken and processed through the S-TRAP protein digest procedure. Ten percent of the eluted peptides were then reconstituted in 0.1% TFA prior to loading onto the timsTOF HT mass spectrometer using the EvoSep One system. Analysis was by a data-dependent acquisition mode with elution from an 8 cm performance column using the 100 SPD method.

The acquired mass spectra were searched against the predicted sequences of proteins encoded by *F. prausnitzii* ATCC27766 draft genome (NCBI accession PXUQ0100), appended with viral protein sequences downloaded from the Uniprot protein resource. A list of common contaminants was added to this database. Searches were carried out using MSFragger version 4.1 running under Fragpipe version 22. The results were then filtered to a target false discovery rate of 1% and a minimum of 2 unique peptides required for identification of a protein (**Table S2**).

Proteins identified by MS analysis but labelled as ‘hypothetical proteins’ were submitted to Collabfold v1.5.5 structure prediction, and Foldseek (GitHub release 8-ef4e960) “easy-search” against the PDB database^76^. Top hits for each hypothetical protein were collated and used to identify other putative functions associated with phage or GTA homologues within loci other than those identified by phage annotation software. Foldseek confidence scores with an E-value cutoff of 0.1 were considered for evaluation.

### Data aggregation and visualisation

Tabular outputs from the above steps were imported into R environment v4.4.0 and processed using a custom script (**Supplementary Dataset** at DOI 10.6084/m9.figshare.26310658). All scaffold properties were summarised in a single table. Taxonomic assignments of prokaryotic scaffolds included in MAGs were based on GTDB-Tk, and on MMseqs2 GTDB search for unbinned scaffolds. Taxonomic assignments of viral scaffolds were derived from ICTV exemplar genome BLASTn search, where possible, and derived from geNomad for the rest of the scaffolds. Based aggregated data, all metagenomic scaffolds were divided into six broad categories: 1) Viruses (VirSorter2 suffix “||full” AND CheckV *Provirus* ≠ “Yes”); 2) MAGs: bacteria (scaffold part of a MAG bin); 3) MAGs: bacteria with prophages (scaffold part of a MAG bin, contains one or multiple sequence fragments of viral origin); 4) Bacteria (unbinned); 5) Bacteria with prophages (unbinned, contains one or multiple sequence fragments of viral origin); 6) Archaea (binned or unbinned).

Relative abundance of a given scaffold was calculated as number of Illumina reads aligned to that scaffold divided by the total number of quality filtered reads for a given sample. Scaffolds enriched in the VLP fractions (**Figure S2**) were identified in the following way. Firstly, a ratio of relative abundance of a given scaffold between a VLP fraction (either top or bottom) and a total community DNA in a given donor was calculated. These ratios were shown to follow normal distribution. Secondly, VLP-overrepresented scaffolds were identified as having their VLP/total community abundance ratios (as Z-scores) above two standard deviations from the mean.

To visualise size distribution and taxonomic origin of individual Nanopore reads, the following filtration steps were applied. Only reads with length ≥ 3kb were considered. Only Minimap2 alignments of ≥ 1kb were kept. In case of ambigous alignment of Nanopore reads to more than one genomic scaffold, a scaffold producing the greatest sum of alignment lengths with a given read was considered as correct template. For plotting purposes, reads aligning to rare biological entities (with ≤ 50 reads per MAG; ≤ 50 reads per unbinned bacterial genomic scaffold; ≤ 200 reads per viral genomic scaffold) were removed.

Package gggenes v0.5.1 was used to visualise DRAM annotated genomic scaffolds, overlayed with Illumina read coverage depth (calculated by Samtools *mpileup* command, smoothed by applying a moving average), and coordinates of Nanopore reads alignments (Minimap2). The rest of the plots were created using ggplot2 v3.5.1 and ggpubr v0.6.0.

## Supplemental material

**Table S1. Packaging events involving bacterial genomic DNA across the three human donors.**

**Table S2. MS analysis of proteins present in a CsCl-purified VLP fraction obtained after spontaneous lysis of a stationary phase *F. prausnitzii* ATCC27766**

**Figure S1. Size distribution and mapping of full-sized nanopore VLP DNA reads obtained from known transducing/packaging phage-bacterium systems. A,** *Bacteroides* phage crAss001. No reads were detected aligning to the genome of the host strain *Bacteroides intestinalis* APC919/174; **B,** *Salmonella* phage P22 infecting its host *S. enterica* LT2, with reads aligning to both the phage and the host genomes (GT); **C,** Enterobacteria phage P1 infecting its host *E. coli* MG1655 (GT); **D,** induction of prophage pp5 in *E. faecalis* VPE14089, ∼44.8 kb-long reads aligning to the prophage region and the adjacent chromosomal region primarily to one side of the prophage, in line with the expected LT pattern^12^; **E,** *B. subtilis* GTA-like element PBSX, ∼13.4 kb-long reads show uniform coverage of the entire *B. subtilis* genome.

**Figure S2. Selection and characteristics of bacterial MAGs enriched in the VLP fractions. A,** relative abundance of genomic scaffolds in the VLP fractions versus the total community fraction deviates from the expected normal distribution (Kolmogorov-Smirnov test); **B,** an arbitrary cut-off of +2 standard deviation (SD) applied to select for genomic scaffolds overrepresented in the VLP fractions; **C,** MAGs containing such overrepresented genomic scaffolds in the top (T) and bottom (B) VLP fractions, dots correspond to individual scaffolds within the MAGs, shaped according to presence (and completeness) of phage sequence in the scaffold, dot colour corresponds to MAG taxonomy, whereas the size reflects scaffold length in bp.

**Figures S3-S8. Length distribution of full-sized nanopore reads produced from DNA extracted from the top (-T) and the bottom (-B) VLP fractions from the three human faecal samples.** Each curve is a separate MAG (top panel)/bacterial species (middle panel)/phage genome (bottom panel); colours represent individual taxa.

**Figure S9. Read coverage of un-binned genomic scaffolds from an uncultured species F23-B02 sp900545805 from the faecal donor 925.** Area plots represent coverage by short Illumina reads (DNA from VLP fractions B and T, as well as the total community DNA), whereas segments of random colours underneath that are full-sized Nanopore reads.

**Figure S10. Read coverage of selected *Bacteroides sp.* genomic scaffolds in the faecal donor 928.** Area plots represent coverage by short Illumina reads (DNA from VLP fractions B and T, as well as the total community DNA), whereas segments of random colours underneath that are full-sized Nanopore reads.

## References

1. Gilbert, J.A., Blaser, M.J., Caporaso, J.G., Jansson, J.K., Lynch, S.V., and Knight, R. (2018). Current understanding of the human microbiome. Nat. Med. 2018 244 24, 392–400. 10.1038/nm.4517.

2. Lozupone, C.A., Stombaugh, J.I., Gordon, J.I., Jansson, J.K., and Knight, R. (2012). Diversity, stability and resilience of the human gut microbiota. Nature 489, 220–230. 10.1038/nature11550.

3. Sommer, F., Anderson, J.M., Bharti, R., Raes, J., and Rosenstiel, P. (2017). The resilience of the intestinal microbiota influences health and disease. Nat. Rev. Microbiol. 15, 630–638. 10.1038/nrmicro.2017.58.

4. Ng, K.M., Aranda-Díaz, A., Tropini, C., Frankel, M.R., Van Treuren, W., O’Loughlin, C.T., Merrill, B.D., Yu, F.B., Pruss, K.M., Oliveira, R.A., et al. (2019). Recovery of the Gut Microbiota after Antibiotics Depends on Host Diet, Community Context, and Environmental Reservoirs. Cell Host Microbe 26, 650–665.e4. 10.1016/j.chom.2019.10.011.

5. Zhu, S., Hong, J., and Wang, T. (2024). Horizontal gene transfer is predicted to overcome the diversity limit of competing microbial species. Nat. Commun. 15, 800. 10.1038/s41467-024-45154-w.

6. Smillie, C.S., Smith, M.B., Friedman, J., Cordero, O.X., David, L.A., and Alm, E.J. (2011). Ecology drives a global network of gene exchange connecting the human microbiome. Nature 480, 241–244. 10.1038/nature10571.

7. Groussin, M., Poyet, M., Sistiaga, A., Kearney, S.M., Moniz, K., Noel, M., Hooker, J., Gibbons, S.M., Segurel, L., Froment, A., et al. (2021). Elevated rates of horizontal gene transfer in the industrialized human microbiome. Cell 184, 2053–2067.e18. 10.1016/J.CELL.2021.02.052.

8. Borodovich, T., Shkoporov, A.N., Ross, R.P., and Hill, C. (2022). Phage-mediated horizontal gene transfer and its implications for the human gut microbiome. Gastroenterol. Rep. 10. 10.1093/GASTRO/GOAC012.

9. Chen, J., Quiles-Puchalt, N., Chiang, Y.N., Bacigalupe, R., Fillol-Salom, A., Chee, M.S.J., Fitzgerald, J.R., and Penadés, J.R. (2018). Genome hypermobility by lateral transduction. Science 362, 207–212. 10.1126/SCIENCE.AAT5867.

10. Humphrey, S., Fillol-Salom, A., Quiles-Puchalt, N., Ibarra-Chávez, R., Haag, A.F., Chen, J., and Penadés, J.R. (2021). Bacterial chromosomal mobility via lateral transduction exceeds that of classical mobile genetic elements. Nat. Commun. 12, 6509. 10.1038/s41467-021-26004-5.

11. Chee, M.S.J., Serrano, E., Chiang, Y.N., Harling-Lee, J., Man, R., Bacigalupe, R., Fitzgerald, J.R., Penadés, J.R., and Chen, J. (2023). Dual pathogenicity island transfer by piggybacking lateral transduction. Cell 186, 3414–3426.e16. 10.1016/J.CELL.2023.07.001.

12. Kleiner, M., Bushnell, B., Sanderson, K.E., Hooper, L.V., and Duerkop, B.A. (2020). Transductomics: sequencing-based detection and analysis of transduced DNA in pure cultures and microbial communities. Microbiome 8, 158. 10.1186/s40168-020-00935-5.

13. Fillol-Salom, A., Bacigalupe, R., Humphrey, S., Chiang, Y.N., Chen, J., and Penadés, J.R. (2021). Lateral transduction is inherent to the life cycle of the archetypical Salmonella phage P22. Nat. Commun. 2021 121 12, 1–12. 10.1038/s41467-021-26520-4.

14. Wolput, S., Lood, C., Fillol-Salom, A., Casters, Y., Albasiony, A., Cenens, W., Vanoirbeek, K., Kerremans, A., Lavigne, R., Penadés, J.R., et al. (2024). Phage-host co-evolution has led to distinct generalized transduction strategies. Nucleic Acids Res. 52, 7780–7791. 10.1093/nar/gkae489.

15. Kogay, R., Koppenhöfer, S., Beatty, J.T., Kuhn, J.H., Lang, A.S., and Zhaxybayeva, O. (2022). Formal recognition and classification of gene transfer agents as viriforms. Virus Evol. 8, veac100. 10.1093/ve/veac100.

16. Kuhn, J.H., and Koonin, E.V. (2023). Viriforms—A New Category of Classifiable Virus-Derived Genetic Elements. Biomolecules 13, 289. 10.3390/biom13020289.

17. Craske, M.W., Wilson, J.S., and Fogg, P.C.M. (2024). Gene transfer agents: structural and functional properties of domesticated viruses. Trends Microbiol. 10.1016/J.TIM.2024.05.002.

18. Sherlock, D., Leong, J.X., and Fogg, P.C.M. (2019). Identification of the First Gene Transfer Agent (GTA) Small Terminase in Rhodobacter capsulatus and Its Role in GTA Production and Packaging of DNA. J. Virol. 93, 10.1128/jvi.01328-19. 10.1128/jvi.01328-19.

19. Tran, N.T., and Le, T.B.K. (2024). Control of a gene transfer agent cluster in Caulobacter crescentus by transcriptional activation and anti-termination. Nat. Commun. 15, 4749. 10.1038/s41467-024-49114-2.

20. Marrs, B. (1974). Genetic Recombination in Rhodopseudomonas capsulata. Proc. Natl. Acad. Sci. 71, 971–973. 10.1073/pnas.71.3.971.

21. Biers, E.J., Wang, K., Pennington, C., Belas, R., Chen, F., and Moran, M.A. (2008). Occurrence and Expression of Gene Transfer Agent Genes in Marine Bacterioplankton. Appl. Environ. Microbiol. 74, 2933–2939. 10.1128/AEM.02129-07.

22. Tomasch, J., Wang, H., Hall, A.T.K., Patzelt, D., Preusse, M., Petersen, J., Brinkmann, H., Bunk, B., Bhuju, S., Jarek, M., et al. (2018). Packaging of Dinoroseobacter shibae DNA into Gene Transfer Agent Particles Is Not Random. Genome Biol. Evol. 10, 359–369. 10.1093/gbe/evy005.

23. Rapp, B.J., and Wall, J.D. (1987). Genetic transfer in Desulfovibrio desulfuricans. Proc. Natl. Acad. Sci. 84, 9128–9130. 10.1073/pnas.84.24.9128.

24. Humphrey, S.B., Stanton, T.B., Jensen, N.S., and Zuerner, R.L. (1997). Purification and characterization of VSH-1, a generalized transducing bacteriophage of Serpulina hyodysenteriae. J. Bacteriol. 179, 323–329. 10.1128/jb.179.2.323-329.1997.

25. Guy, L., Nystedt, B., Toft, C., Zaremba-Niedzwiedzka, K., Berglund, E.C., Granberg, F., Näslund, K., Eriksson, A.S., and Andersson, S.G.E. (2013). A Gene Transfer Agent and a Dynamic Repertoire of Secretion Systems Hold the Keys to the Explosive Radiation of the Emerging Pathogen Bartonella. PLOS Genet. 9, e1003393. 10.1371/JOURNAL.PGEN.1003393.

26. Gozzi, K., Tran, N.T., Modell, J.W., Le, T.B.K., and Laub, M.T. (2022). Prophage-like gene transfer agents promote Caulobacter crescentus survival and DNA repair during stationary phase. PLOS Biol. 20, e3001790. 10.1371/journal.pbio.3001790.

27. Bertani, G. (1999). Transduction-like gene transfer in the methanogen methanococcus voltae. J. Bacteriol. 181, 2992–3002. 10.1128/JB.181.10.2992-3002.1999.

28. Modi, S.R., Lee, H.H., Spina, C.S., and Collins, J.J. (2013). Antibiotic treatment expands the resistance reservoir and ecological network of the phage metagenome. Nature 499, 219–222. 10.1038/nature12212.

29. Enault, F., Briet, A., Bouteille, L., Roux, S., Sullivan, M.B., and Petit, M.A. (2016). Phages rarely encode antibiotic resistance genes: a cautionary tale for virome analyses. ISME J. 2017 111 11, 237–247. 10.1038/ismej.2016.90.

30. Lücking, D., Mercier, C., Alarcón-Schumacher, T., and Erdmann, S. (2023). Extracellular vesicles are the main contributor to the non-viral protected extracellular sequence space. ISME Commun. 3, 112. 10.1038/s43705-023-00317-6.

31. Chiura, H.X., Kogure, K., Hagemann, S., Ellinger, A., and Velimirov, B. (2011). Evidence for particle-induced horizontal gene transfer and serial transduction between bacteria. FEMS Microbiol. Ecol. 76, 576–591. 10.1111/J.1574-6941.2011.01077.X.

32. Domingues, S., and Nielsen, K.M. (2017). Membrane vesicles and horizontal gene transfer in prokaryotes. Curr. Opin. Microbiol. 38, 16–21. 10.1016/J.MIB.2017.03.012.

33. Casjens, S.R., and Gilcrease, E.B. (2009). Determining DNA Packaging Strategy by Analysis of the Termini of the Chromosomes in Tailed-Bacteriophage Virions. Methods Mol. Biol. Clifton NJ 502, 91–111. 10.1007/978-1-60327-565-1_7.

34. Rao, V.B., and Feiss, M. (2008). The Bacteriophage DNA Packaging Motor. Annu. Rev. Genet. 42, 647–681. 10.1146/annurev.genet.42.110807.091545.

35. Bárdy, P., Füzik, T., Hrebík, D., Pantůček, R., Thomas Beatty, J., and Plevka, P. (2020). Structure and mechanism of DNA delivery of a gene transfer agent. Nat. Commun. 2020 111 11, 1–13. 10.1038/s41467-020-16669-9.

36. Shkoporov, A.N., Khokhlova, E.V., Fitzgerald, C.B., Stockdale, S.R., Draper, L.A., Ross, R.P., and Hill, C. (2018). ΦCrAss001 represents the most abundant bacteriophage family in the human gut and infects Bacteroides intestinalis. Nat. Commun. 9, 4781. 10.1038/s41467-018-07225-7.

37. Casjens, S., and Hayden, M. (1988). Analysis in vivo of the bacteriophage P22 headful nuclease. J. Mol. Biol. 199, 467–474. 10.1016/0022-2836(88)90618-3.

38. Sternberg, N. (1990). Bacteriophage P1 cloning system for the isolation, amplification, and recovery of DNA fragments as large as 100 kilobase pairs. Proc. Natl. Acad. Sci. U. S. A. 87, 103. 10.1073/pnas.87.1.103.

39. Matos, R.C., Lapaque, N., Rigottier-Gois, L., Debarbieux, L., Meylheuc, T., Gonzalez-Zorn, B., Repoila, F., Lopes, M. de F., and Serror, P. (2013). Enterococcus faecalis Prophage Dynamics and Contributions to Pathogenic Traits. PLOS Genet. 9, e1003539. 10.1371/journal.pgen.1003539.

40. Andersont, L.M., and Bott, K.F. (1985). DNA packaging by the Bacillus subtilis defective bacteriophage PBSX. J. Virol. 54, 773–780. 10.1128/JVI.54.3.773-780.1985.

41. Dutilh, B.E., Cassman, N., McNair, K., Sanchez, S.E., Silva, G.G.Z., Boling, L., Barr, J.J., Speth, D.R., Seguritan, V., Aziz, R.K., et al. (2014). A highly abundant bacteriophage discovered in the unknown sequences of human faecal metagenomes. Nat. Commun. 5, 4498. 10.1038/ncomms5498.

42. Cornuault, J.K., Petit, M.A., Mariadassou, M., Benevides, L., Moncaut, E., Langella, P., Sokol, H., and De Paepe, M. (2018). Phages infecting Faecalibacterium prausnitzii belong to novel viral genera that help to decipher intestinal viromes. Microbiome 6, 65. 10.1186/S40168-018-0452-1.

43. Frazão, N., Konrad, A., Amicone, M., Seixas, E., Güleresi, D., Lässig, M., and Gordo, I. (2022). Two modes of evolution shape bacterial strain diversity in the mammalian gut for thousands of generations. Nat. Commun. 2022 131 13, 1–14. 10.1038/s41467-022-33412-8.

44. Frazão, N., Sousa, A., Lässig, M., and Gordo, I. (2019). Horizontal gene transfer overrides mutation in *Escherichia coli* colonizing the mammalian gut. Proc. Natl. Acad. Sci. 116, 17906–17915. 10.1073/pnas.1906958116.

45. Wang, S., Jiang, Y., Che, L., Wang, R.H., and Li, S.C. (2024). Enhancing insights into diseases through horizontal gene transfer event detection from gut microbiome. Nucleic Acids Res. 52, e61. 10.1093/nar/gkae515.

46. Coyne, M.J., Zitomersky, N.L., McGuire, A.M., Earl, A.M., and Comstock, L.E. (2014). Evidence of Extensive DNA Transfer between *Bacteroidales* Species within the Human Gut. mBio 5. 10.1128/mBio.01305-14.

47. Dahlman, S., Avellaneda-Franco, L., Kett, C., Subedi, D., Young, R.B., Gould, J.A., Rutten, E.L., Gulliver, E.L., Turkington, C.J.R., Nezam-Abadi, N., et al. (2023). Temperate gut phages are prevalent, diverse, and predominantly inactive. Preprint at bioRxiv, 10.1101/2023.08.17.553642 https://doi.org/10.1101/2023.08.17.553642.

48. Kim, M.-S., and Bae, J.-W. (2018). Lysogeny is prevalent and widely distributed in the murine gut microbiota. ISME J. 12, 1127–1141. 10.1038/s41396-018-0061-9.

49. Reyes, A., Haynes, M., Hanson, N., Angly, F.E., Heath, A.C., Rohwer, F., and Gordon, J.I. (2010). Viruses in the faecal microbiota of monozygotic twins and their mothers. Nature 466, 334–338. 10.1038/nature09199.

50. Avellaneda-Franco, L., Dahlman, S., and Barr, J.J. (2023). The gut virome and the relevance of temperate phages in human health. Front. Cell. Infect. Microbiol. 13. 10.3389/fcimb.2023.1241058.

51. Hu, J., Ye, H., Wang, S., Wang, J., and Han, D. (2021). Prophage Activation in the Intestine: Insights Into Functions and Possible Applications. Front. Microbiol. 12, 785634. 10.3389/FMICB.2021.785634.

52. Oh, J.-H., Alexander, L.M., Pan, M., Schueler, K.L., Keller, M.P., Attie, A.D., Walter, J., and van Pijkeren, J.-P. (2019). Dietary Fructose and Microbiota-Derived Short-Chain Fatty Acids Promote Bacteriophage Production in the Gut Symbiont *Lactobacillus reuteri*. Cell Host Microbe 25, 273–284.e6. 10.1016/j.chom.2018.11.016.

53. Henrot, C., and Petit, M.A. (2022). Signals triggering prophage induction in the gut microbiota. Mol. Microbiol. 118, 494–502. 10.1111/MMI.14983.

54. Arumugam, M., Raes, J., Pelletier, E., Le Paslier, D., Yamada, T., Mende, D.R., Fernandes, G.R., Tap, J., Bruls, T., Batto, J.-M., et al. (2011). Enterotypes of the human gut microbiome. Nature 473, 174–180. 10.1038/nature09944.

55. Penadés, J.R., Chen, J., Quiles-Puchalt, N., Carpena, N., and Novick, R.P. (2015). Bacteriophage-mediated spread of bacterial virulence genes. Curr. Opin. Microbiol. 23, 171–178. 10.1016/j.mib.2014.11.019.

56. Touchon, M., Moura de Sousa, J.A., and Rocha, E.P. (2017). Embracing the enemy: the diversification of microbial gene repertoires by phage-mediated horizontal gene transfer. Curr. Opin. Microbiol. 38, 66–73. 10.1016/j.mib.2017.04.010.

57. Fillol-Salom, A., Alsaadi, A., de Sousa, J.A.M., Zhong, L., Foster, K.R., Rocha, E.P.C., Penadés, J.R., Ingmer, H., and Haaber, J. (2019). Bacteriophages benefit from generalized transduction. PLOS Pathog. 15, e1007888. 10.1371/JOURNAL.PPAT.1007888.

58. Haaber, J., Leisner, J.J., Cohn, M.T., Catalan-Moreno, A., Nielsen, J.B., Westh, H., Penadés, J.R., and Ingmer, H. (2016). Bacterial viruses enable their host to acquire antibiotic resistance genes from neighbouring cells. Nat. Commun. 2016 71 7, 1–8. 10.1038/ncomms13333.

59. Lang, A.S., Zhaxybayeva, O., and Beatty, J.T. (2012). Gene transfer agents: phage-like elements of genetic exchange. Nat. Rev. Microbiol. 10, 472–482. 10.1038/nrmicro2802.

60. Lang, A.S., and Beatty, J.T. (2007). Importance of widespread gene transfer agent genes in α-proteobacteria. Trends Microbiol. 15, 54–62. 10.1016/j.tim.2006.12.001.

61. Banks, E.J., and Le, T.B.K. (2024). Co-opting bacterial viruses for DNA exchange: structure and regulation of gene transfer agents. Curr. Opin. Microbiol. 78, 102431. 10.1016/j.mib.2024.102431.

62. Québatte, M., and Dehio, C. (2019). Bartonella gene transfer agent: Evolution, function, and proposed role in host adaptation. Cell. Microbiol. 21, e13068. 10.1111/cmi.13068.

63. Eiserling, F., Pushkin, A., Gingery, M., and Bertani, G. (1999). Bacteriophage-like particles associated with the gene transfer agent of methanococcus voltae PS. J. Gen. Virol. 80 (Pt 12), 3305–3308. 10.1099/0022-1317-80-12-3305.

64. Wood, H.E., Dawson, M.T., Devine, K.M., and McConnell, D.J. (1990). Characterization of PBSX, a defective prophage of Bacillus subtilis. J. Bacteriol. 172, 2667–2674. 10.1128/jb.172.5.2667-2674.1990.

65. Krogh, S., O’Reilly, M., Nolan, N., and Devine, K.M. (1996). The phage-like element PBSX and part of the skin element, which are resident at different locations on the Bacillus subtilis chromosome, are highly homologous. Microbiology 142, 2031–2040. 10.1099/13500872-142-8-2031.

66. Fitzgerald, C.B., Shkoporov, A.N., Sutton, T.D.S., Chaplin, A.V., Velayudhan, V., Ross, R.P., and Hill, C. (2018). Comparative analysis of Faecalibacterium prausnitzii genomes shows a high level of genome plasticity and warrants separation into new species-level taxa. BMC Genomics 19, 1–20. 10.1186/S12864-018-5313-6.

67. Johnson, C.M., and Grossman, A.D. (2015). Integrative and Conjugative Elements (ICEs): What They Do and How They Work. Annu. Rev. Genet. 49, 577–601. 10.1146/annurev-genet-112414-055018.

68. Mandelbaum, N., Zhang, L., Carasso, S., Ziv, T., Lifshiz-Simon, S., Davidovich, I., Luz, I., Berinstein, E., Gefen, T., Cooks, T., et al. (2023). Extracellular vesicles of the Gram-positive gut symbiont Bifidobacterium longum induce immune-modulatory, anti-inflammatory effects. NPJ Biofilms Microbiomes 9, 30. 10.1038/s41522-023-00400-9.

69. Liu, H., Tian, Y., Xue, C., Niu, Q., Chen, C., and Yan, X. (2022). Analysis of extracellular vesicle DNA at the single-vesicle level by nano-flow cytometry. J. Extracell. Vesicles 11, e12206. 10.1002/jev2.12206.

70. Faddetta, T., Vassallo, A., Del Duca, S., Gallo, G., Fani, R., and Puglia, A.M. (2022). Unravelling the DNA sequences carried by Streptomyces coelicolor membrane vesicles. Sci. Rep. 2022 121 12, 1–8. 10.1038/s41598-022-21002-z.

71. Biller, S.J., Mcdaniel, L.D., Breitbart, M., Rogers, E., Paul, J.H., and Chisholm, S.W. (2016). Membrane vesicles in sea water: heterogeneous DNA content and implications for viral abundance estimates. ISME J. 2017 112 11, 394–404. 10.1038/ismej.2016.134.

72. Camargo, A.P., Roux, S., Schulz, F., Babinski, M., Xu, Y., Hu, B., Chain, P.S.G., Nayfach, S., and Kyrpides, N.C. (2024). Identification of mobile genetic elements with geNomad. Nat. Biotechnol. 42, 1303–1312. 10.1038/s41587-023-01953-y.

73. Zhou, Y., Liang, Y., Lynch, K.H., Dennis, J.J., and Wishart, D.S. (2011). PHAST: a fast phage search tool. Nucleic Acids Res. 39, W347–352. 10.1093/nar/gkr485.

74. Arndt, D., Grant, J.R., Marcu, A., Sajed, T., Pon, A., Liang, Y., and Wishart, D.S. (2016). PHASTER: a better, faster version of the PHAST phage search tool. Nucleic Acids Res. 44, W16–21. 10.1093/nar/gkw387.

75. Guo, J., Bolduc, B., Zayed, A.A., Varsani, A., Dominguez-Huerta, G., Delmont, T.O., Pratama, A.A., Gazitúa, M.C., Vik, D., Sullivan, M.B., et al. (2021). VirSorter2: a multi-classifier, expert-guided approach to detect diverse DNA and RNA viruses. Microbiome 9, 37. 10.1186/s40168-020-00990-y.

76. van Kempen, M., Kim, S.S., Tumescheit, C., Mirdita, M., Lee, J., Gilchrist, C.L.M., Söding, J., and Steinegger, M. (2024). Fast and accurate protein structure search with Foldseek. Nat. Biotechnol. 42, 243–246. 10.1038/s41587-023-01773-0.

